# Microrobots Powered by Concentration Polarization Electrophoresis (CPEP)

**DOI:** 10.1101/2022.10.14.512287

**Authors:** Florian Katzmeier, Friedrich C. Simmel

## Abstract

Second-order electrokinetic flow around colloidal particles caused by concentration polarization electro-osmosis (CPEO) can result in a phoretic motion of asymmetric particle dimers in a homogeneous AC electrical field, which we refer to as concentration polarization electro-phoresis (CPEP). To demonstrate this actuation mechanism, we created particle dimers from micron-sized silica spheres with sizes 1.0 µm and 2.1 µm by connecting them with DNA linker molecules. The dimers can be steered along arbitrarily chosen paths within a 2D plane by controlling the orientation of the AC electric field in a fluidic chamber with the joystick of a gamepad. Further utilizing induced dipole-dipole interactions, we demonstrate that particle dimers can be used to controllably pick up monomeric particles and release them at any desired position, and also to assemble several particles into groups. Systematic experiments exploring the dependence of the dimer migration speed on the electric field strength, frequency, and buffer composition align with the theoretical framework of CPEO and provide parameter ranges for the operation of our microrobots.

---

According to Smoluchowski’s century-old theory, electrophoresis of colloidal particles is shape-independent [1]. In combination with the time reversibility of hydrodynamics at low Reynolds numbers, shape-independence implies that even asymmetric particles will not display any net movement when subjected to a homogeneous AC electric field. However, under experimental conditions which generate nonlinear electrokinetic phenomena, particles with a broken symmetry can experience directed propulsion also in homogeneous AC electric fields. Such AC electrophoretic propulsion was first theoretically proposed [2] for *strongly* polarizable particles based on induced charge electroosmosis (ICEO) [3] and later experimentally verified for metallo-dielectric Janus particles, which were observed to move perpendicular to the electric field direction [4].

In this work, we investigate a novel propulsion mechanism for *weakly* polarizable particles with a non-zero surface charge based on the phenomenon of concentration polarization electroosmosis (CPEO). CPEO was recently theoretically described [5] and experimentally validated [5–8] and is found to produce similar flow patterns around spheres in an AC electric field as ICEO, but under different experimental conditions. We therefore expected that similar to propulsion via induced charge electrophoresis (ICEP) resulting from ICEO, asymmetric particles subjected to CPEO would also experience directed propulsion, which we accordingly refer to as concentration polarization electrophoresis (CPEP).

In the most widely utilized experimental setup, microswimmers are placed on an electrode and exposed to a vertical electric field. Within this setup, it was demonstrated that asymmetric colloidal dimers [9] and metallo-dielectric Janus particles [10] are propelled perpendicularly to the electric field in a random direction in the 2D plane. To introduce maneuverability, magnetic fields have been used in combination with ferromagnetic metallo-dielectric Janus particles [11] and ferromagnetic asymmetric colloidal dimers [12]. Further, it has been demonstrated that metallodielectric Janus particles can be used to transport other dielectric particles [11, 13, 14]. In the case of asymmetric colloidal dimers, the propulsion mechanism is based on the electrohydrodynamic interplay between electrode and particles [15, 16].

Since, in contrast to this propulsion mechanism, CPEO does not require an electrode in close proximity, we surmised that it could be applied to propel asymmetric colloidal dimer particles using an *in-plane* electric field. Taking advantage of electro-orientation, which orients prolate particles parallel to an AC electric field through induced dipole alignment and induced hydrodynamic flows [17–20], we thus expected to achieve directed propulsion of asymmetric dimers *along* the field lines rather than perpendicular to them.

In the following, we demonstrate that asymmetric dimer ‘microrobots’ can be precisely maneuvered using a straightforward electrical setup without any additional magnetic forces by simply controlling the orientation of an homogeneous AC electric field in the plane of movement. Such an AC electric-controlled 2D actuation was previously only achieved through dielectrophoresis [21], which requires electric field gradients and a computer-controlled feedback mechanism [18, 19]. We also develop a strategy to pick up, transport, and release spherical cargo particles with these microrobots by making use of induced dipole-dipole interactions and hydrodynamic flow fields. Finally, we explore the dependence of the microrobots’ migration speeds on electric field strength, frequency, and buffer composition, which is found to agree reasonably well with the theoretical predictions of CPEO.

## RESULTS & DISCUSSION

### Asymmetric colloidal microswimmers in an AC electrical field

It is known that the axisymmetric fluid flow depicted in Fig. 1b arises around weakly polarizable particles with a non-zero surface charge when subjected to an AC electrical field in a low-ionic strength aqueous medium. Fluid flows towards the particle in the direction of the electric field and is repelled perpendicularly to the electric field [5, 7]. We expected that for an asymmetric dimeric particle an asymmetric flow would arise as proposed in Fig. 1c that would lead to the propulsion of the particle. Further, a dimeric particle will also align with the external electric field as shown in Fig. 1c and Fig. 1a due to an induced electric dipole. In combination with the propulsion this leads to a directed motion along the field lines of the electric field. The movement of the dimers can thus be easily controlled by changing the direction and strength of the external AC electric field.

**FIG. 1.**
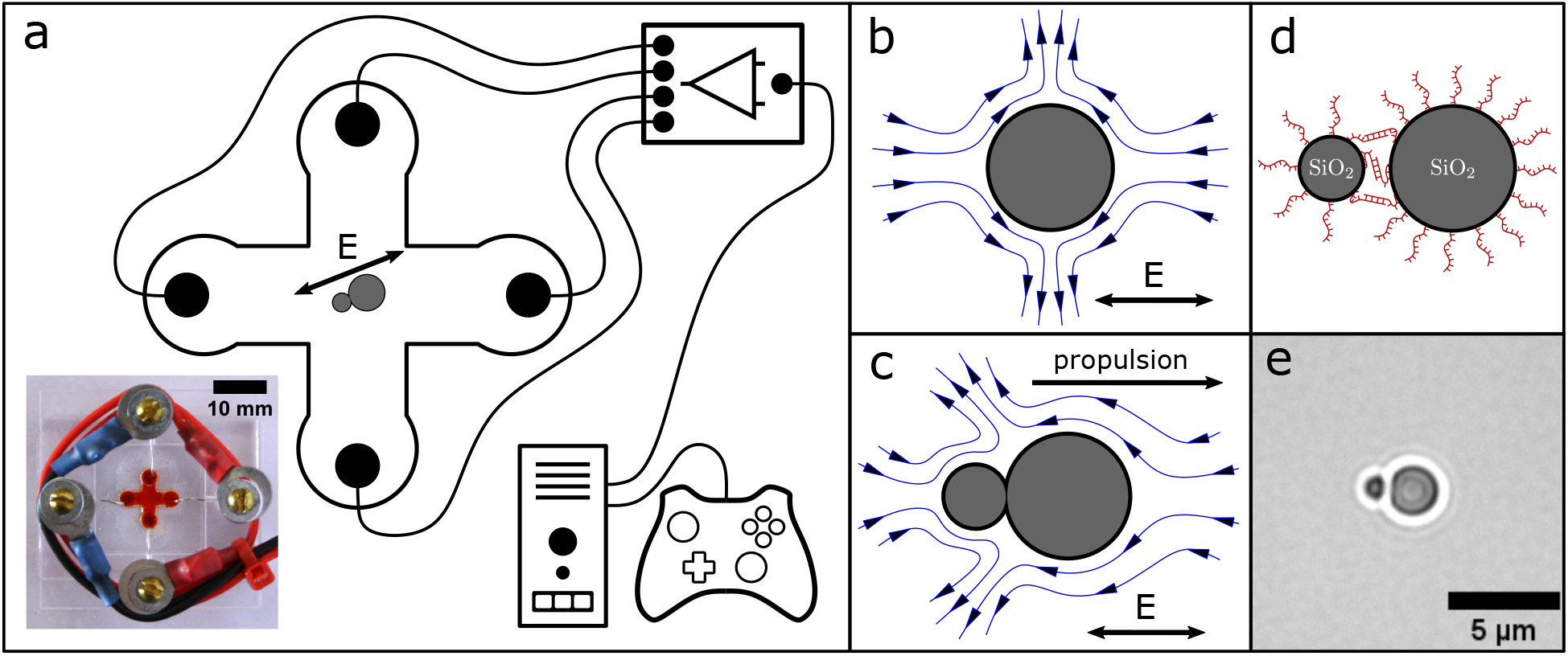
(a) Scheme and photograph of our experimental setup that enables control of the direction and amplitude of an AC electric field in a microscopy chamber. A dimer is drawn in the center of the cross-shaped fluidic chamber, which aligns with the externally applied AC field through an induced dipole. For visualization, the fluidic chamber in the photograph is filled with a red dye. (b) Electrokinetic flow around a spherical particle arising in an AC electric field. (c) Expected electrokinetic flow around an asymmetric particle dimer in an AC electric field, which results in directed propulsion. (d) DNA modified colloids form a dimer through DNA hybridization. (d) Microscopy image of a particle dimer.

### Experimental setup and fabrication of particle dimers

For our experiments we designed the sample chamber shown in Fig. 1a, in which two microfluidic channels equipped with platinum electrode pairs at their inlets intersect in the center. The electric field in the center of the chamber is a superposition of the fields generated by the remote electrode pairs. Hence, the field at the intersection is homogeneous, and its direction and amplitude can be controlled by applying different electric field strengths to the channels [22, 23]. We created two electric signals with the sound card of a computer, which were amplified in two stages using custom-built amplifiers before feeding them into the microchannels. With our setup we can apply AC voltages with an amplitude of up to 305 V which corresponds to an electric field amplitude of approximately 60 mV*/*µm in the center of our chamber. We programmed a python script to control the amplitude of the electric signals via the XY-deflection of the analog joystick of a gamepad (an Xbox Controller) which is conventionally used to play video games. As a result, the direction and amplitude of the AC electric field in our sample chamber and thereby the movement of our microrobots can be directly controlled with a joystick while imaging them with a microscope. We also included the possibility to change the field frequency to predefined values 250 Hz and 750 Hz by pressing the buttons available on the controller. Images of the setup together with detailed information on its design and manufacture can be found in the Supplementary Materials.

Asymmetric particle dimers acting as microrobots were synthesized through the self-assembly of two differently sized, DNA-coated silica spheres with diameters 1.01 µm and 2.12 µm, respectively [24–27]. To this end, each particle type was modified with 60 nt long single-stranded DNA molecules, which had 30 nt long sub-sequences that were complementary to sequences on the respective other particle type. When mixed in the presence of 4 mM MgCl_2_, the silica spheres specifically bound to each other via DNA duplex formation (cf. Fig. 1d & e). For our experiments, we diluted the dimers in Tris buffer (100 µM, pH 8.4) supplemented with 5.2 µM MgCl_2_. A detailed description of synthesis and sample preparation is given in the Methods.

### Movement and maneuverability of the microrobots

Our protocol for the assembly of the silica particles resulted in a mixture of mainly monomers and dimers with only small amounts of higher oder multimers. Upon exposure to an AC electric field in our sample chamber, the dimers are subject to a torque due to an induced dipole which aligns the dimer axis in parallel to the electric field lines. The dimers can assume two alternative, stable orientations in the AC field, in which the positions of the larger and smaller particle are exchanged with each other (Fig. 2a). Notably, the induced hydrodynamic flow around each dimer is asymmetric and propels it in the direction defined by the larger particle. Thus the particle dimers shown in the scheme of Fig. 2a would be expected to move in opposite directions, as indicated by the blue pointers.

**FIG. 2.**
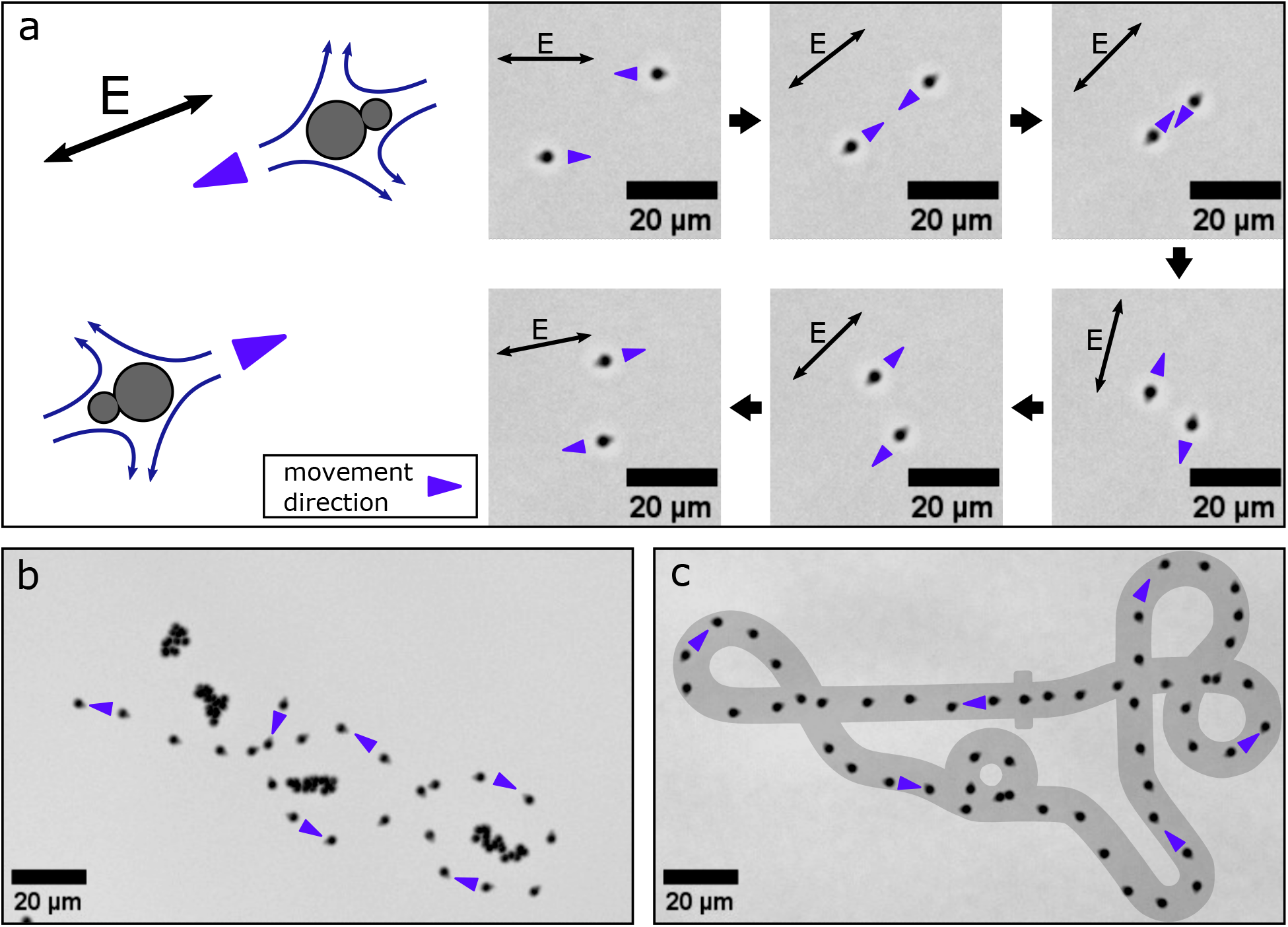
Controlled movement of particle dimers. (a) The sketch on the left shows two microrobots aligned with an AC electric field. The orientation of the is indicated with a black double arrow. The movement direction is indicated with a blue pointer and is opposite for the two microrobots due to their opposite orientations. The image sequence on the right shows successive frames of a microscopy video demonstrating the resulting anti-synchronous movement in an electric field with slowly changing orientation. (b) By applying joystick-controlled AC voltages a microrobot is maneuvered through a slalom course around stationary monomer particles. The image shows an overlay of successive frames of a recorded microscopy video. Due to Brownian motion, the monomer particles appear as particle clouds, but they do not respond the applied AC field. (c) A microrobot is maneuvered along a race track adopted from a well-known video game. The image shows an overlay of successive frames of a recorded microscopy video and the racetrack.

To demonstrate this effect in the experiment, we recorded a microscopy video of two differently aligned dimers while slowly changing the direction of the applied electric field using the joystick. As expected, the dimers were observed to move anti-synchronously, meaning that the trajectory of one particle was the point reflection of the other (the image sequence shown in Fig. 2a is the first part of Supplementary Video 1; details on video processing are given in the Methods section). All dimers in a sample move along the electric field lines collectively, with the larger particles in the front. We thus focused on the movement of individual dimers in all further experiments.

To demonstrate microrobot maneuverability, we recorded a microscopy video, in which we steered a microrobot along a slalom course around islands of monomeric particles, which remained stationary in the AC field (cf. Figure 2b and second part of Supplementary Video 1). The monomeric particles appear as clouds in the overlay image, since they are subject to Brownian motion. We also found a slight drift in our microscopy videos due to bulk fluid motion which we corrected by tracking several of the stationary monomeric particles and shifting the recorded video by their average displacement.

We also recorded a microscopy video, in which we maneuvered a microrobot along a racetrack adopted from a computer game. For this purpose, we printed the racetrack on a cling film and attached it with tape to the screen of the computer controlling the microscope to enable visual feedback and control by a human operator. Figure 2c shows an overlay of the racetrack and video images recorded during the experiment (cf. second part of Supplementary Video 1).

### Pick-up, transport, and release of cargo particles

We found that the microrobots can be readily used to pick up, transport, and release other, *monomeric* cargo particles, for which we utilized both electric and hydrodynamic interactions between microrobot and cargo. Figure 3a shows a microscopy image of a microrobot approaching a cargo particle (shown in green in the image). As illustrated in the sketch above the microscopy image, fluid flows towards both particles in the direction of the electric field and is repelled perpendicularly to it. The microrobot and the cargo particle drift in the fluid flow caused by each other which results in an attractive interaction for the configuration shown.

**FIG. 3.**
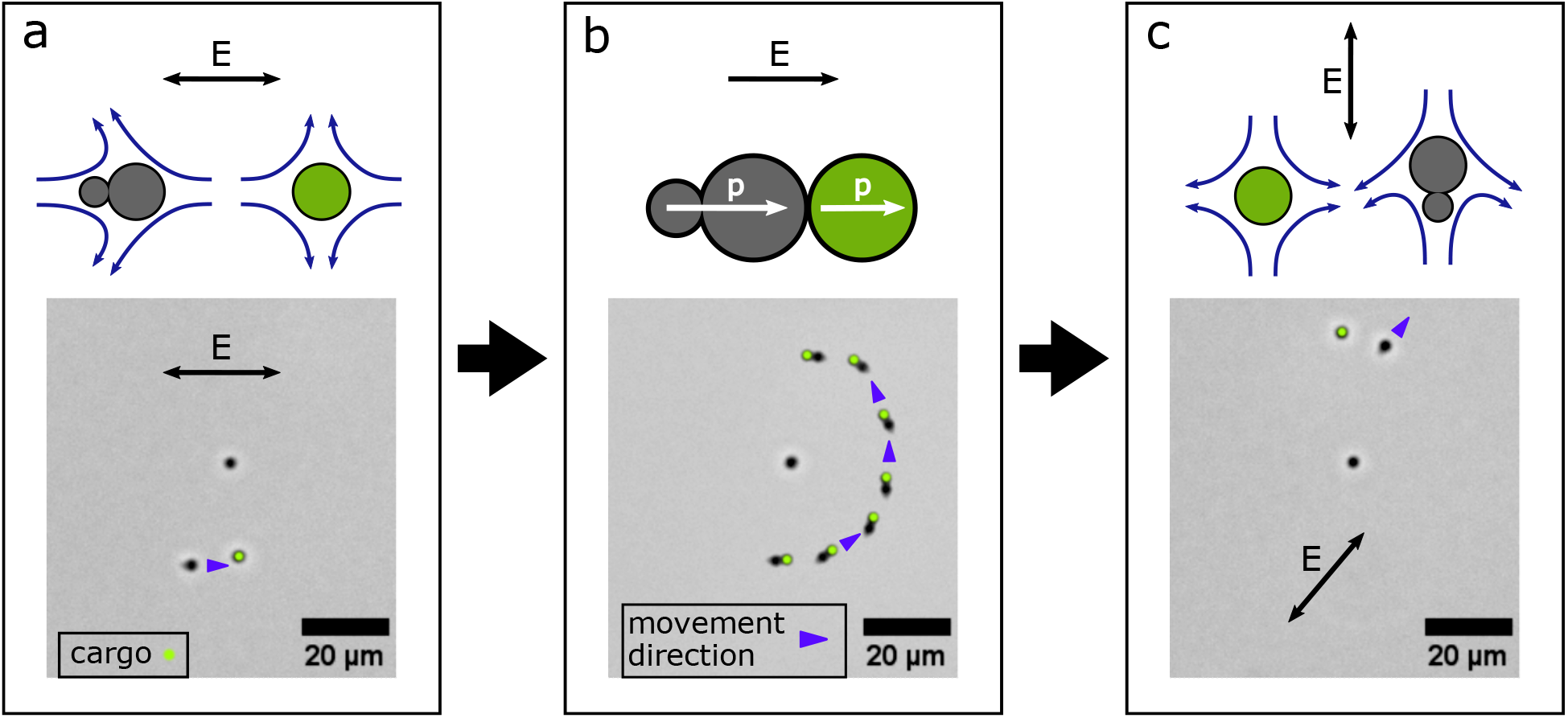
Cargo pick-up, transport and release. (a) A microrobot approaches a cargo particle. The direction of motion of the microrobot is indicated with blue pointer and the cargo particle is labeled with a green dot. The orientation of the AC electric field is indicated with a black double arrow. The fluid flow arising around the microrobot and the cargo particle is illustrated with curved blue arrows in the sketch above and leads to an attraction. (b) A microrobot sticks to a cargo particle via induced dipole-dipole forces. Both are maneuvered around another monomeric particle. The instantaneous induced dipole moments are indicated with white arrows in the sketch above, the electric field is indicated with a black arrow. (c) A cargo particle is released from a microrobot by a quick change in direction of the external electric field. The fluid flow arising around the microrobot and the cargo particle now leads to repulsion.

When in direct contact, the microrobot sticks to the cargo via induced dipole-dipole forces (see Fig. 3b). Even though we apply an AC electric field, at any point in time the external field induces electric dipoles in both microrobot and cargo, which point in the same direction and thus result in a near-field attraction of the particles. This mechanism is well known and results in particle chain formation in crowded colloidal suspensions. [7, 28, 29]. We then maneuvered the cargo-loaded microrobot around another monomeric particle as shown in the overlay image in Fig. 3b. For this image, we corrected the drift in the corresponding microscopy video by moving the tracked position of the monomeric *non-cargo* particle into the center of each frame.

Cargo release (shown in Fig. 3c) was achieved by switching off the electric field, changing the frequency from 250 Hz to 750 Hz, and then applying an AC electric field with an orientation roughly perpendicular to the previous field. The corresponding particle and field configuration is illustrated in the sketch above the microscopy image: the fluid is repelled perpendicularly to the electric field from the particles’ equators, resulting in repulsion of the particles. As soon as the microrobot had moved sufficiently far away from the cargo particle, the frequency was set back to 250 Hz, which re-established electrokinetic control over the microrobot. We found that changing the frequency made it easier for the experimenter to execute cargo release. We assume that at higher frequencies the induced dipole-dipole force decreases in strength relative to the force exerted by the fluid flow. A video of the cargo transport process is shown in the fourth part of Supplementary Video 1.

### Controlled assembly of cargo particles into particle chains

Using the same strategy for particle transport and release, we were also able to assemble several monomeric particles into a particle chain. As shown in Figure 4 (cf. last part of Supplementary Video 1), the microrobot can be controlled to sequentially pick up two individual cargo particles and drop them off in the vicinity of a third target particle. As a result of the attractive induced dipole-dipole interactions between the monomeric particles, the three particles stick together and form a particle chain.

**FIG. 4.**
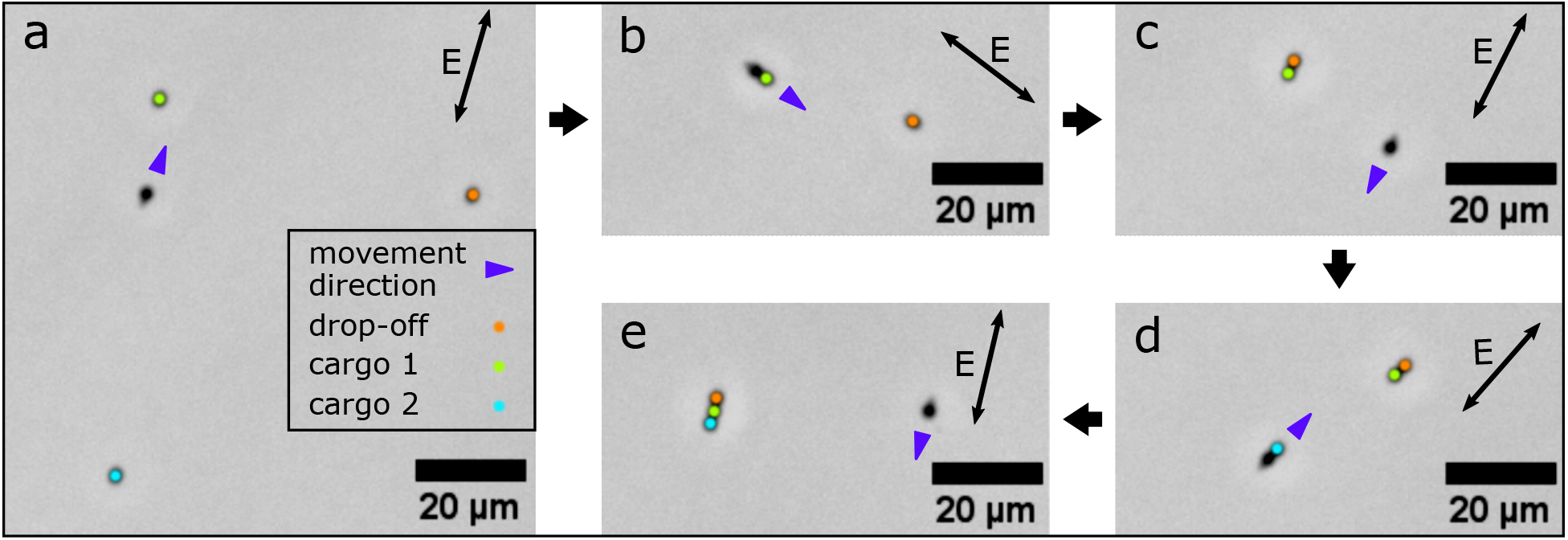
Assembly of three monomeric particles into a particle chain. (a) A microrobot approaches cargo particle 1 (labeled with a green dot). (b) The microrobot, loaded with cargo 1, approaches another monomeric particle (orange), which serves as the drop-off location for cargo release. (c) The cargo particle sticks to the orange particle via induced dipole-dipole forces, while the microrobot is maneuvered towards cargo particle 2 (turquoise). (d) The microrobot loaded with cargo 2 heads back towards the drop-off location to release the cargo. (e) The two cargo particles and the orange target particle are assembled into a chain.

### Amplitude and frequency dependence of the CPEO mechanism

Having established a novel approach for the manipulation of asymmetric colloidal dimers, we intended to verify whether the underlying propulsion mechanism indeed conformed with the theoretical framework for CPEO. We hypothesized that the migration speed of the dimers would scale similarly as the strength of the fluid flow around spherical monomer particles. Like ICEO, CPEO is a second-order phenomenon with respect to the applied electric field and thus the migration speed *v* should scale with the electric field amplitude *E*_0_ as 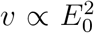. However, the frequency dependence of CPEO is expected to differ substantially from that of ICEO. Notably, the fluid velocity around spherical particles caused by CPEO falls off to zero for frequencies exceeding the characteristic frequency 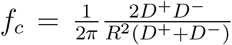 [5, 7, 30, 31]. Here, *D*^*−*^ and *D*^+^ denote the diffusion coefficients of the buffer ions Cl^−^ and Tris – H^+^, which are *D*^*−*^ = 2 *×* 10^3^ µm^2^*/*s and *D*^+^ = 0.8 *×* 10^3^ µm^2^*/*s at *T* = 20 ^*°*^C [32] and *R* is the radius of the object. When applying the theory to our dimers, we can interpret *R* as their typical size, which we take as the average radius of their constituent particles, and with the given parameters we obtain *f*_*c*_ = 297 Hz. 1*/*(2*πf*_*c*_) corresponds to the time required for ions to diffuse over the distance *R*. By contrast, in the case of ICEO the characteristic frequency is derived from the time required to charge the electric double layer on the particle, which is given by the RC time of the corresponding circuit [3, 33].

We measured the migration speeds of three different dimers for several voltages and frequencies to verify the above hypothesis. To this end, we employed the setup shown in Fig. 5a where dimers are placed in a linear microchannel with two electrodes at its opposite inlets. As before, we prepared the dimers in 100 µM Tris-buffer, which we titrated to pH 8.4 by the addition of HCl and supplemented with 5.2 µM MgCl_2_. We then recorded microscopy videos of the migration of the three dimers while applying AC fields with different frequencies and amplitudes. From the start and end positions of the dimers, we computed the distances covered and from these the average migration speeds. We also measured the speed of a monomeric particle as a reference. The corresponding measurements are listed in Tables S9-S11. Fig. 5c shows plots of the migration speeds versus the applied electric field amplitude at a constant frequency of 250 Hz. As shown, the experimental migration speeds are well described by a quadratic fit 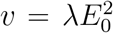. In Fig. 5b, the frequency dependence of the migration speeds of the three dimers is plotted for a constant electric field amplitude of 16.8 mV*/*µm. We find a decrease of the migration speed in the range of the characteristic frequency *f*_*c*_ = 297 Hz calculated for CPEO flows.

**FIG. 5.**
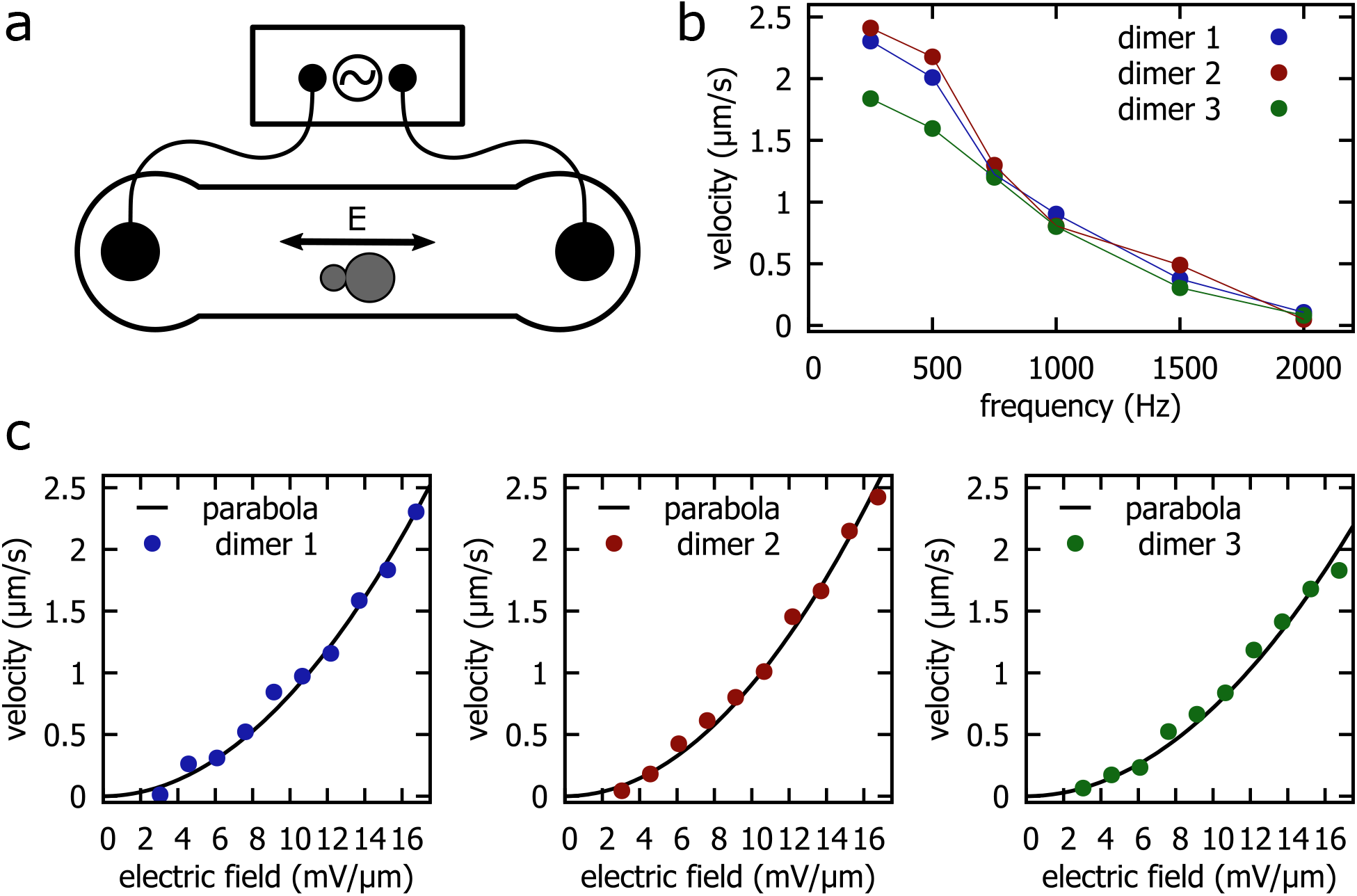
Dependence of migration speed on frequency and amplitude of the AC field. (a) Simplified measurement setup. An AC electric field is applied to a linear microscopy channel containing dimers. (b) Migration speed of three different dimers at a constant electric field amplitude of 16.8 mV*/*µm and varying frequency. The colored lines are a guide for the eye. (c) Migration speed of three different dimers at a constant frequency of 250 Hz and varying electric field amplitude. The black lines are parabolas 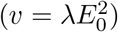 fitted to the velocity data.

For comparison, we computed the characteristic frequency of ICEO flows around metal and uncharged dielectric spheres at our experimental conditions [3, 33]. In addition, we calculated the expected slip velocities around spheres for CPEO and ICEO flows at the highest applied electric field amplitude 16.8 mV*/*µm (Table I) [3, 5, 34]. The corresponding calculation can be found in the Supplementary Information. As mentioned, the frequency response of our dimers agrees best with the characteristic frequency predicted by CPEO, whereas *f*_*c*_ predicted by ICEO for dielectric particles is two orders of magnitude off. The characteristic frequency predicted for strongly polarizable particles, such as metal particles, is closer to the experimentally observed value, but application of this model to our case is physically unreasonable as silica particles are not strongly polarizable. The value for the slip velocity around a sphere calculated from CPEO is found to be one order of magnitude larger than the observed migration speed of our dimers. This result is not unexpected since the slip velocity and the migration speed are not directly equivalent, as also demonstrated in the schematic diagram shown in Fig 1c. The migration speed may be further reduced due to the additional drag caused by the nearby channel bottom. Importantly, the slip velocity calculated for ICEO flow around a dielectric sphere is approximately one order of magnitude lower than the observed dimer migration speed.

**TABLE I.**
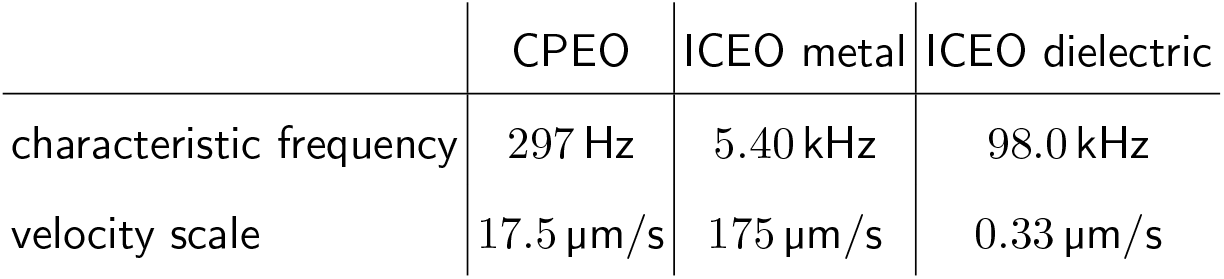
Characteristic frequencies *f*_*c*_ and velocity scales *v*

### Buffer dependence of the transport mechanism

We finally also characterized the buffer dependence of the microrobots’ migration velocity. CPEO flows are caused by ion-selective surface conduction in the electric double layer at the particle surface, which depends on the zeta potential. The zeta potential, in turn, is a function of the surface charge of the particle and the ionic strength of the buffer solution. The flow is thus expected to be strongest for large surface potentials, i.e., under conditions with large surface charge densities and low ionic strengths. We recorded microscopy videos of dimers prepared in buffers with different concentrations of Tris, NaCl and NaOH, each supplemented with 5.2 µM MgCl_2_. In addition, we tested Tris-buffer, NaCl and NaOH without any MgCl_2_ and also a solution containing exclusively MgCl_2_. For each buffer composition, we recorded tracks of at least five dimers, and we took care that every video contained at least one spherical particle as a reference. As before, we measured the migration speed by marking the start and end position of the dimer and dividing the resulting distance by the elapsed time. We also measured the migration speed of all spherical particles in each microscopy video and used it as a reference (see Methods for details). The results of these experiments are listed in Tables S1-S8 and plotted in Fig. 6.

**FIG. 6.**
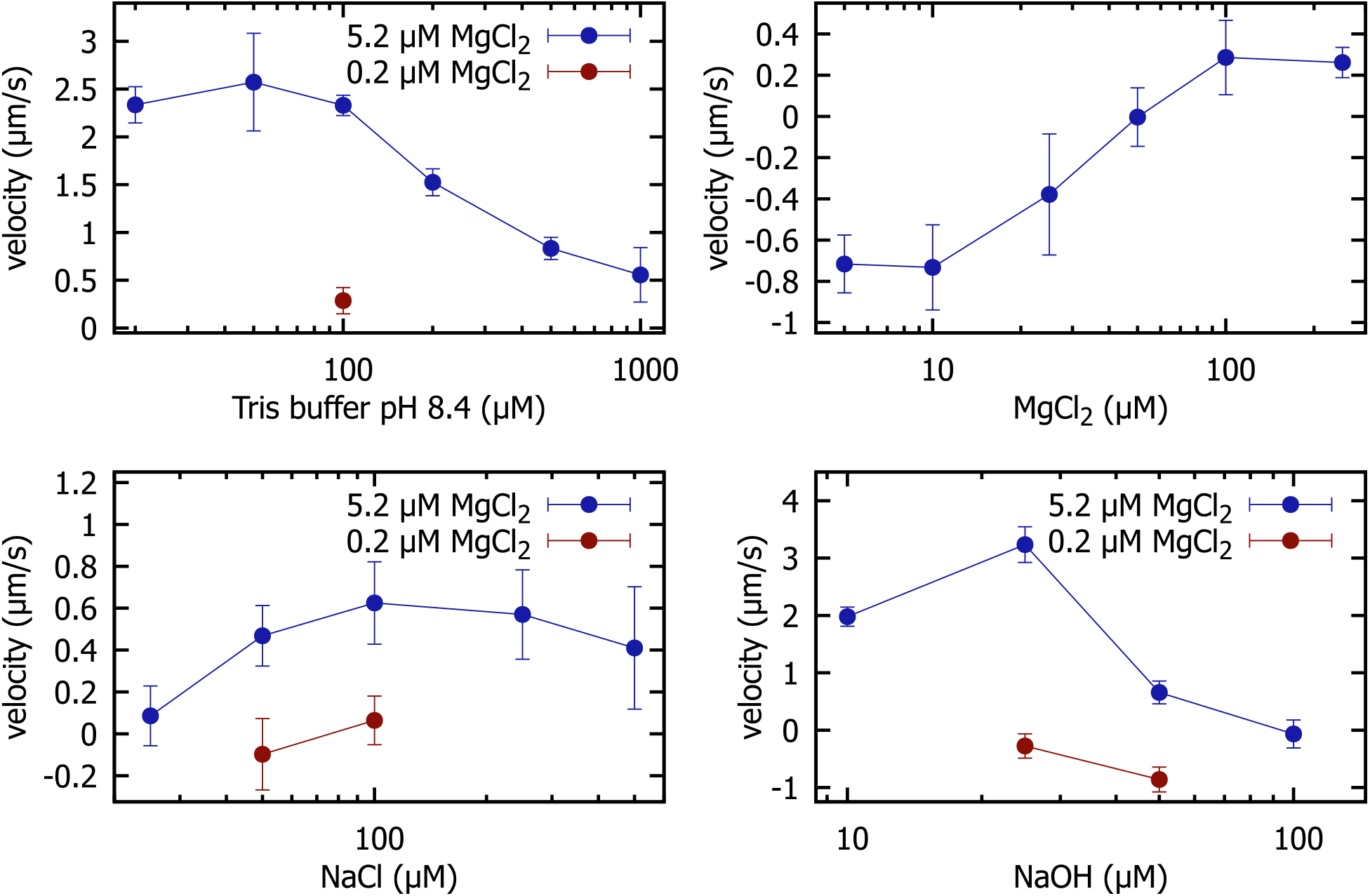
Dimer velocities at a constant electric field amplitude of 16.8 mV*/*µm and constant frequency of 250 Hz for various buffer conditions. Blue data points indicate measurements with MgCl_2_ supplemented to the buffer, while red data points correspond to measurements without supplemented MgCl_2_. Each data point is the average of at least 5 velocity measurements of different dimers, error bars indicate the corresponding standard deviations. Data points with negative velocities correspond to dimers moving backward, which we defined as a movement with the smaller particle in front.

Overall, the velocities tended to decrease for increasing monovalent salt concentrations, approaching zero velocity for concentrations around 1 mM, which is in the range expected for CPEO flows for typical values of the surface charge [7]. The details of the buffer dependence of the particle velocity are intricate, however. We found that supplementing the buffers with 5.2 µM MgCl_2_ had a tremendous effect on the migration behavior. For Tris buffer, we found a strong enhancement of the velocity by MgCl_2_. For NaOH, we found that dimers moved backwards in the absence of MgCl_2_, while they moved forward in its presence. Notably, when using MgCl_2_ in dH_2_O only, we found backward movement that changed to forward movement at higher concentrations. When present alone and at low concentrations, either NaOH or MgCl_2_ induced backward movement. However, when combined they induced forward movement. We found the largest migration velocities for NaOH and Tris buffer at pH 8.4, which we attribute to an increase in surface charge caused by the elevated pH. As even small amounts of MgCl_2_ had an extreme effect on the migration velocity, we took special care to avoid any salt contamination in our samples (see Methods for details).

A complex dependence on buffer conditions for an electrokinetic phenomenon such as CPEO it is not entirely unexpected. Our initial picture for fluid flow around a particle dimer (Fig. 1c), was motivated by the expectation that larger flows will occur around the larger particle in the dimer, which sets the propulsion direction. Strictly speaking, we have no reason to assume a specific propulsion direction as no theory is currently available for CPEO flows around asymmetric particles. Interestingly, we find that the theory of Fernández-Mateo et al. [5], although only valid for binary electrolytes with equal ion diffusion coefficients *D*, already predicts flow reversal around spherical particles for certain values of *D*. We plotted their result for the flow magnitudes for varying values of *D* in the Supplementary Information. A detailed mechanistic explanation of the observed buffer dependence is further complicated by the surface modification of the silica particles with DNA, which is known to have a strong affinity for Mg^2+^ ions [35].

We would finally like to comment on the propulsion mechanism for metallo-dielectric Janus particles and colloidal dimers exposed to a vertical electric field. For metallo-dielectric Janus particles, a CPEO flow may occur in addition to the ICEO flow generated on the metallic side, when the dielectric Janus face is sufficiently charged. As ICEO and CPEO flows have different characteristic frequencies, the interplay of these effects will depend on the applied frequency. A similar case can be made for particle dimers exposed to a vertical electric field, which should also be subject to CPEO flows when applying AC electric fields in the frequency range set by the corresponding characteristic frequency.

## CONCLUSION

We have introduced a novel approach towards AC electrophoretic manipulation of colloidal microswimmers, which facilitates precise electrical control over two-dimensional movements. In contrast to other approaches for electrically driven swimmers, the utilization of concentration–polarization electroosmosis (CPEO) and electro-orientation enables the use of in-plane electric fields to move the particles in the direction of the field lines (rather than perpendicular to them as in other approaches). The directed movement requires asymmetric particles with a surface charge, but the particles themselves do not have to be ‘Janus’ or magnetic, which broadens the design possibilities for electrically controlled microrobots. In our case, two differently sized silica particles were connected using DNA linker molecules. We employed a relatively simple setup to achieve 2D actuation where we need no additional magnetic fields, as in the case of dimers subject to a vertical electric field, nor computer-controlled feedback, as in the case of dielectrophoretic-driven microswimmers. As indicated by the joystick-controlled actions demonstrated in this work, our approach is of immediate interest for applications in microrobotics. The microrobots can move along arbitrarily chosen paths in 2D and can be directed to controllably pick up, release, and also assemble particles into groups.

Finally, we confirmed that the dependence of our microrobots’ migration speed on AC field frequency, amplitude, and electrolyte concentration aligns with the theoretical expectations for CPEO-driven movement. In future work, AC electrophoretic manipulation could be used to implement more complex robotic functions. For instance, differently sized/shaped particles are expected to show a different frequency response, which could also be used to frequency-select only a fraction of the microrobots. It is conceivable to let microrobots assemble other microparticles into defined superstructures, potentially these assembled structures could also act as microrobots themselves. One of the main challenges for future applications is the realization of operating conditions that are compatible with useful chemical or biochemical reactions (to assist assembly tasks), or even with biological entities, e.g., for the manipulation of bacteria or even mammalian cells.

## METHODS

### Functionalization and dimerization of colloidal particles

Carboxylated silica spheres with diameters 1.01 µm (Lot: SiO_2_ – COOH-AR756-5ml) and 2.12 µm (Lot: SiO_2_ – COOH-AR1060-5ml) were purchased from microParticles GmbH. We modified the surface of the silica spheres by activating the carboxyl groups with 1-Ethyl-3-(3-dimethylaminopropyl) carbodiimide (EDC) and coupling them to amino-modified DNA. [36, 37] The colloids were reacted in 200 µL of 100 mM MES buffer (pH 4.8 adjusted with HCl and NaOH) containing 250 µM amino-modified DNA and 250 mM EDC (Merck: Art. No. E6383-1G) on a rotator at room temperature for 3 h. We used colloid concentrations of 11.35 · 10^9^/mL and 50 · 10^9^/mL of the 2.12 µm colloids and the 1.01 µm colloids, respectively, to account for the different surface areas of the colloids. The colloids were then washed and incubated extensively in borate buffer (boric acid adjusted to pH 8.2 with NaOH) and deionized water to get rid of remaining reaction components and to hydrolyze unreacted activated carboxyl groups. We avoided using buffers containing amino groups for washing as we wanted to preserve the negative surface charge of the colloids. An extended protocol with details on the washing procedure is given in the Supplementary Materials. Finally, the colloids were diluted to concentrations of 2.27 · 10^9^/mL (2.12 µm colloids) and 10 · 10^9^/mL (1.01 µm colloids) in deionized water, shock frozen in liquid nitrogen and stored at *−*80 ^*°*^C.

Our two DNA strands are 60 nucleotides (nt) long and are each composed of a 30 nt long spacer region followed by a 30 nt region which is complementary to the corresponding region on the other strand. The spacer provides flexibility in the distance between the colloids where hybridization can take place. We designed our DNA sequences with NUPACK [38] such that they have no secondary structure. The oligonucleotides were purchased from Integrated DNA Technologies as dried pellets, their sequences are listed in the Supplementary information. We diluted our DNA strands in deionized water and stored them at *−*20 ^*°*^C.

### Microrobot assembly

We assembled our microrobots by incubating concentrations of approximately 1.6 · 10^9^/mL of each colloid with 4 mM MgCl_2_ in a reaction volume of 25 µL for 45 min on a rotator. The above colloid concentration assumes that no colloids were lost in the above washing procedure. The reaction is stopped by rapid dilution of the sample by a factor of 1 to 1000 in deionized water. The sample is handled with special care as we found that shaking causes the microrobots to disintegrate. For further use, we usually diluted our microrobots again by a factor of 1 to 20 in a buffer of choice. The microrobots were assembled freshly for every day of experiments.

### Sample preparation

In initial experiments, we found a reduction in the migration speed after washing our pipette tips. We therefore suspected that the pipette tips contained trace amounts of divalent ions. In order to establish stable and reproducible behavior of the swimmers, we henceforth cleaned all pipette tips and the sample chamber with deionized water before usage. With every fresh pipette tip, we pipetted deionized water three times before pipetting an actual sample. We also washed all used reaction tubes with deionized water before usage, vortexed them and removed the deionized water again. For our screening experiments, we used commercial microscopy chambers purchased from ibidi (µ-Slide VI 0.4; Cat.No:80601). Before usage, the microscopy chambers were filled three times with deionized water and then blown dry with nitrogen gas. For our experiments with Tris-buffer, we created a 50 mM stock solution at pH 8.4 by titrating Tris (Carl Roth: Art. No. 4855.2) with HCl. We avoided using NaOH in case of overshooting pH 8.4 as this would have resulted in an unknown concentration of additional NaCl in the buffer.

### Video editing

We used ImageJ to edit our videos. We corrected the drift in the corresponding microscopy video by tracking the monomeric particles with the ImageJ plugin TrackMate [39] and shifting the recorded video by their displacement. Overlay images were created by computing the minimum intensity of a collection of frames from a microscopy video. We adjusted the contrast and brightness of our videos and images such that they appear alike. The final video editing was done with the freely available software Shotcut [40].

### Data analysis

We marked the start and end positions of every microrobot and reference particle in a video and saved the coordinates with the corresponding frame number. We also recorded the instantaneous orientations of every microrobot, which lets us identify backward and forward movements. For that purpose, we used an ImageJ macro to automatize the data analysis. We computed the velocities of all microrobots and reference particles by subtracting the y-coordinates of the start and end positions and dividing the result by the elapsed time. The elapsed time was extracted from the metadata of the corresponding video. We then computed the average velocity of all reference particles in a video and subtracted the velocity of every microrobot in a video by the result, which gives us the corrected microrobot velocities. The average and standard deviation are then computed from the corrected velocities of all videos with the same buffer conditions. All measured velocities and the recorded microrobot orientations for our buffer characterization experiments are listed in the Supplementary Tables S1-S8. In our frequency and electric field characterization experiments, we were interested in the response of a single microrobot and reference particle. We therefore applied a simplified data analysis procedure and measured only the speed of the single microrobot and reference particle in a video. The recorded measurements are listed in the Supplementary Tables S9-S11.

### Setup design and operation

The design of our self-made sample chamber is inspired by that of Kopperger et al. [22]. Our sample chambers were milled from of a 5 mm thick PMMA sheet using a micro-milling machine. The bottom part of our microscopy chamber is a glass cover slide (Carl Roth: Art. No. CEX2.1). The PMMA part is glued to the glass slide with Dichloromethane (Carl Roth: Art. No. 8424.2). We use platinum wires (Merck: Art. No. 267201-400MG) with a diameter of 0.5 mm as electrodes. The electrode mounting is milled from a 10 mm PMMA sheet. We use a standard Xbox Controller (PDP 049-012-EU-RD Controller Xbox Series X Rot) connected via an USB-cable to the microscope computer to control electrical signals generated by the sound card of the computer. A more detailed description, including photographs of the setup, can be found in the Supplementary Materials. For microscopy, we used an Olympus IX71 inverted microscope equipped with a 20x objective (Olympus UPlanFL N 20x/0.50) and an 100x objective (Olympus PlanApo 100x/1.40 Oil). During our screening experiments we monitored the current and voltage with a digital oscilloscope (PicoScope 2000) to avoid systematic errors. Electric signals for our screening experiments were created with a function generator (RIGOL DG812) and amplified with an amplifier built in-house.

### Programming

We used the online tool ChatGPT [41] based on GPT3 [42], a Generative Pretrained Transformer developed by OpenAI, to assist with programming.

## SUPPLEMENTARY INFORMATION

The Supplementary Information contains a description of the design of the experimental setup and its operation; calculations of the characteristic frequencies and velocity scales; a discussion of the flow reversal phenomenon; a detailed protocol for the functionalization of the colloids; tables with raw data; supplementary references. The Supporting Video 1 contains videos corresponding to the images shown in Figures 2-4.

## ACKNOWLEDGMENTS

We thank Thomas Mayer for helping us with the DNA strand design. We thank Jonathan List for providing the second stage of our custom-built amplifiers.

## DECLARATIONS

### Ethics approval

Not applicable

### Availability of data and materials

All data needed to evaluate the conclusions in the paper are present in the paper or the Supplementary Materials. Raw video data are available by the authors upon request.

### Code availability

The program code to control our microrobots and the data analysis scripts are available by the authors upon reasonable request.

### Author contributions

F.K. and F.C.S. conceived the project. F.K. planned the experiments, built the experimental setup, performed all experiments, and analyzed the data. F.K. wrote the paper with support by F.C.S.

