## Supplementary Information for "Microrobots Powered by Concentration Polarization Electrophoresis (CPEP)"

(Dated: March 9, 2023)

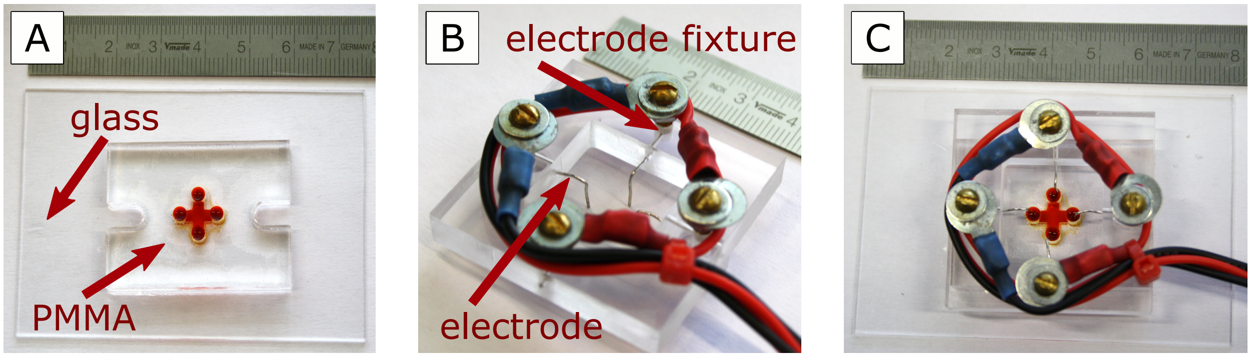


FIG. 1. Photographs of our setup. (A) Sample chamber filled with a red dye. (B) Electrode mounting. (C) Electrode mounting placed on the sample chamber.

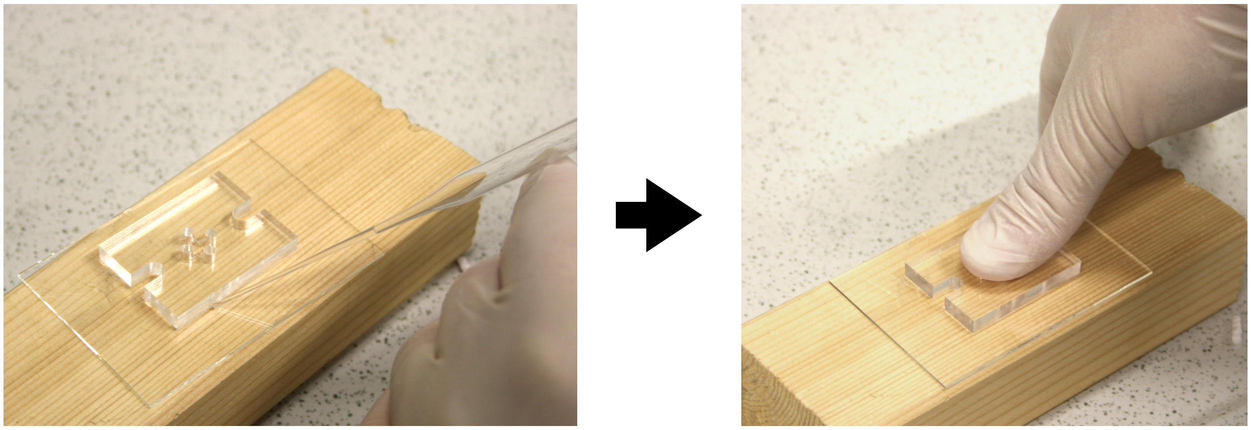

FIG. 2. Illustration of the glueing process used for manufacturing the sample chamber.

### I. DESIGN OF THE EXPERIMENTAL SETUP AND ITS OPERATION

Photographs of our experimental setup are shown in Fig. S1. The sample chamber is shown in Fig. S1A, where it is filled with a red dye for better visualization. The sample chamber consists of two parts. Its bottom is constituted by a glass cover slide, whereas its top is made from a piece of PMMA, from which the channel geometry has been milled out via micro-milling. Both parts are glued together with dichloromethane. The gluing process is illustrated in Fig. S2. First, the PMMA part is placed on the glass cover slide. Then small drops of dichloromethane are placed with a glass pipette at the edge of the PMMA part. Due to capillary forces, the drops wet the surface between the PMMA and the glass slide. The dichloromethane dissolves the surface of the PMMA. The PMMA part is then pressed gently with a finger onto the glass cover slide

until the dichloromethane is dried. Fig. S1B and Fig. S2C show the electrode mounting, where four electrodes are fixed on a PMMA frame. The electrodes extend out of small holes from the PMMA frame and are clamped with a screw from above (see red arrow in Fig. S1B with the label "electrode fixture"). In Fig. S2C the electrode mounting is placed on the sample chamber such that the electrodes extend into the inlets of the sample chamber, resulting in an operational setup.

### II. CHARACTERISTIC FREQUENCIES AND VELOCITY SCALES

In this section, we compute the characteristic frequencies and magnitudes of the slip velocity for CPEO flows around charge dielectric particles and for ICEO flow around dielectric and metallic particles. We use the average particle radius $R = 0.78\,\mu\text{m}$ of the two particles composing our dimers as the typical size. Further, we use the inverse averaged diffusion constant as used by [1] and [2] for the diffusion constant which is defined by $D = \frac{2D^+D^-}{D^++D^-}$. $D^+$ and $D^-$ are the ion diffusion constants for TrisH$^+$ and Cl$^-$, respectively, and are given by $D^+ = 800\,\mu\text{m}^2/\text{s}$ and $D^- = 2000\,\mu\text{m}^2/\text{s}$. Numerically, $D = 1143\,\mu\text{m}^2/\text{s}$ for this case.

The characteristic frequency of ICEO flows around metal particles is derived from the charging time $\tau = \lambda R/D$[3] of the electric double layer on the particle surface, where $\lambda$ is the Debye screening length and depends on the ionic strength $c$. A concentration of $c = 50\,\mu\text{M}$ corresponds to $\lambda = 43\,\text{nm}$. The characteristic frequency is then given by

$$f_c = \frac{1}{2\pi}\frac{D}{\lambda R}, \tag{1}$$

which results in $f_c = 5396$ Hz.

For ICEO flows around dielectric particles the characteristic time scale is given by $\tau = \lambda^2/D$ [3] which corresponds to a characteristic frequency of

$$f_c = \frac{1}{2\pi}\frac{D}{\lambda^2}, \tag{2}$$

which gives $f_c = 98.0\,\text{kHz}$.

Note that the equation for the characteristic time scale for dielectric particles in [3] has an additional erroneous factor $\epsilon_w/\epsilon_d$ (see their equation 6.14). $\epsilon_w/\epsilon_d$ is the fraction of the dielectric constants of water and the particle. The equation above can be obtained when starting from their equation 6.7 and by following the steps stated in the paper. Additionally, one has to assume $\frac{1+\epsilon_w/\epsilon_d}{\epsilon_w/\epsilon_d} \approx 1$, which is reasonable as in most cases $\epsilon_w \gg \epsilon_d$.

The characteristic time scale for CPEO flows is independent of the Debye screening length and is given by $\tau = R^2/D$[1]. $\tau$ is here the time scale ions need to diffuse across the typical length scale of our dimer. The characteristic frequency is given by

$$f_c = \frac{1}{2\pi}\frac{D}{R^2}, \tag{3}$$

which for our parameter settings is $f_c = 297\,\text{Hz}$.

The slip velocity $V$ for ICEO flows around a metal particle is given by

$$V = \epsilon_w \frac{RE^2}{\mu} \tag{4}$$

where $\epsilon_w$ is the dielectric constant of water and $\mu$ is the viscosity. When using the maximum applied electric field amplitude $E = 16.75\,\text{mV}/\mu\text{m}$, we obtain $V = 0.33\,\mu\text{m/s}$.

The slip velocity $V$ for ICEO flows around a dielectric particles is given by

$$V = \frac{3}{4}\epsilon_d \frac{\lambda E^2}{\mu} \tag{5}$$

where $\epsilon_d$ is the dielectric constant of the particle [4]. We used the relative permittivity of 3.7 for silica to compute $\epsilon_d$. When again using the maximum applied electric field amplitude we get $V = 0.33\,\mu\text{m/s}$.

The slip velocity $V$ for CPEO flows around a charged dielectric particle is given by

$$V = U\epsilon_w \frac{RE^2}{\mu} \tag{6}$$

where $U$ is the dimensionless velocity computed which has typically a magnitude of $U = 0.1$ (see below and in [1]). For the maximum applied electric field amplitude this results in $V = 17.47\,\mu\text{m/s}$.

### III. FLOW REVERSAL

In this section, we plot the result of [1] for the dimensionless flow magnitude $U$ of CPEO flows around spherical particles for different ion diffusion constants. Their result for $U$ is a function of the zeta potential $\zeta$, the Dukhin number $Du$, and the ion diffusion constant $D$. The Dukhin number is a dimensionless quantity that characterizes the surface conduction of colloidal particles. In Fig. 3, we plot the dimensionless flow magnitude $U$ versus the dimensionless frequency $2\pi f/(D/R^2)$ for different ion diffusion constants $D$ and Dukhin numbers $Du$ at a constant zeta potential $\zeta = 101\,\mathrm{mV}$. The plot on the upper left is a recreation of a plot shown in [1], where we plotted the frequency response of $U$ for the ion diffusion constants of KCl ($D = 2036\,\mathrm{\mu m^2/s}$) and four different values of the Dukhin number. In the other plots, we decreased the diffusion constant. We find decreasing velocities for the higher Dukhin numbers, 1 and 10, for decreasing values of the ion diffusion constant. For very low diffusion constants ($D = 226\,\mathrm{\mu m^2/s}$) and Dukhin numbers of 1 and 10, we find flow reversal at low frequencies.

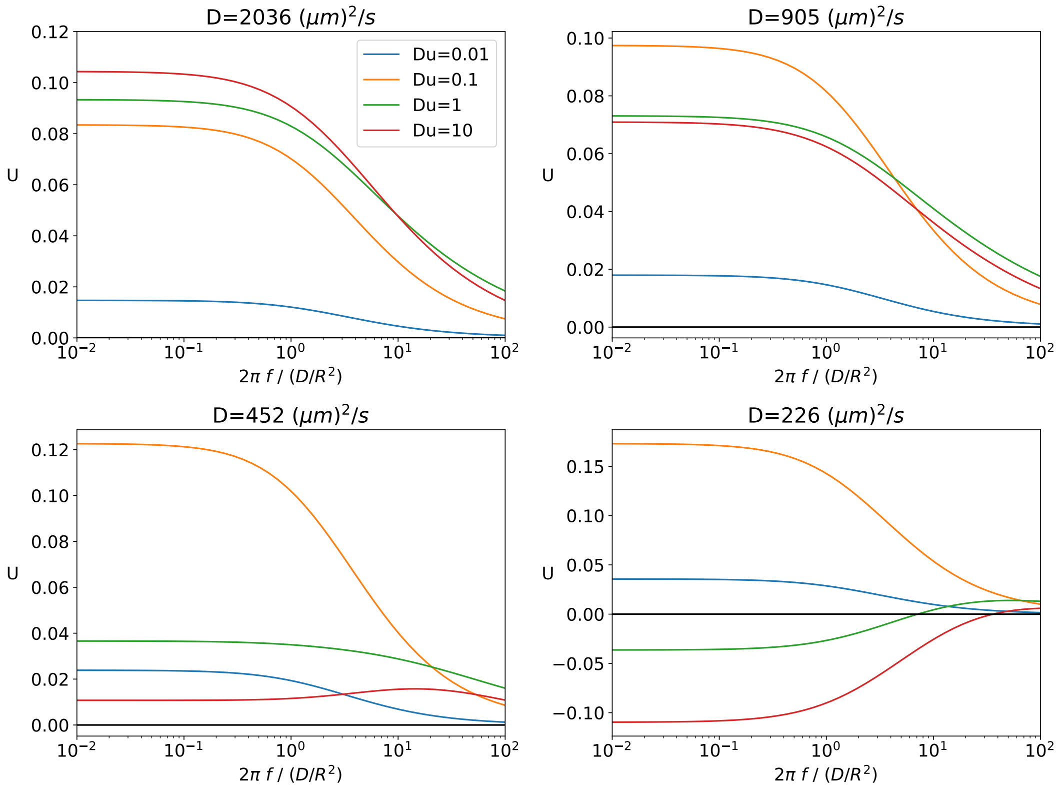


FIG. 3. Plots of the frequency response of the dimensionless flow magnitude of CPEO flows around spheres for different diffusion constants $D$ and Dukhin numbers $Du$. The dimensionless zeta potential was set to $\zeta = -4$. The upper left plot is a recreation of the plot shown in [1]

In Fig. 4, we systematically plotted the frequency response of the flow magnitude for different values of the surface charge density $\sigma$. Within one graph, we plotted the frequency response for several values of the ion diffusion constant. Note that the zeta potential $\zeta$ is a function of the surface charge density and the Debye length $\lambda$, which is in turn a function of the salt concentration $c$ that we set to $\lambda = 43\,\text{nm}$ and $c = 50\,\mu\text{M}$. We assume a particle radius of $R = 0.78\,\mu\text{m}$. We vary the surface charge since both the Dukhin number and the zeta potential are functions of the surface charge.For low surface charges ($\sigma = -0.0005\,\text{C/m}^2$, $\sigma = -0.0010\,\text{C/m}^2$, $\sigma = -0.0015\,\text{C/m}^2$) and low zeta potentials, we find overall increasing flow magnitudes $U$ for increasing surface charges. However, at high surface charges ($\sigma = -0.0045\,\text{C/m}^2$, $\sigma = -0.0080\,\text{C/m}^2$, $\sigma = -0.0160\,\text{C/m}^2$), we find decreasing flow magnitudes for increasing surface charges, which decrease more strongly for lower ion diffusion constants. This leads, at a certain point, to a flow reversal at low frequencies. Interestingly, the graphs cross the zero line at higher frequencies and become positive again. Overall, we find that flow reversal can be found for high surface charges and low diffusion constants.

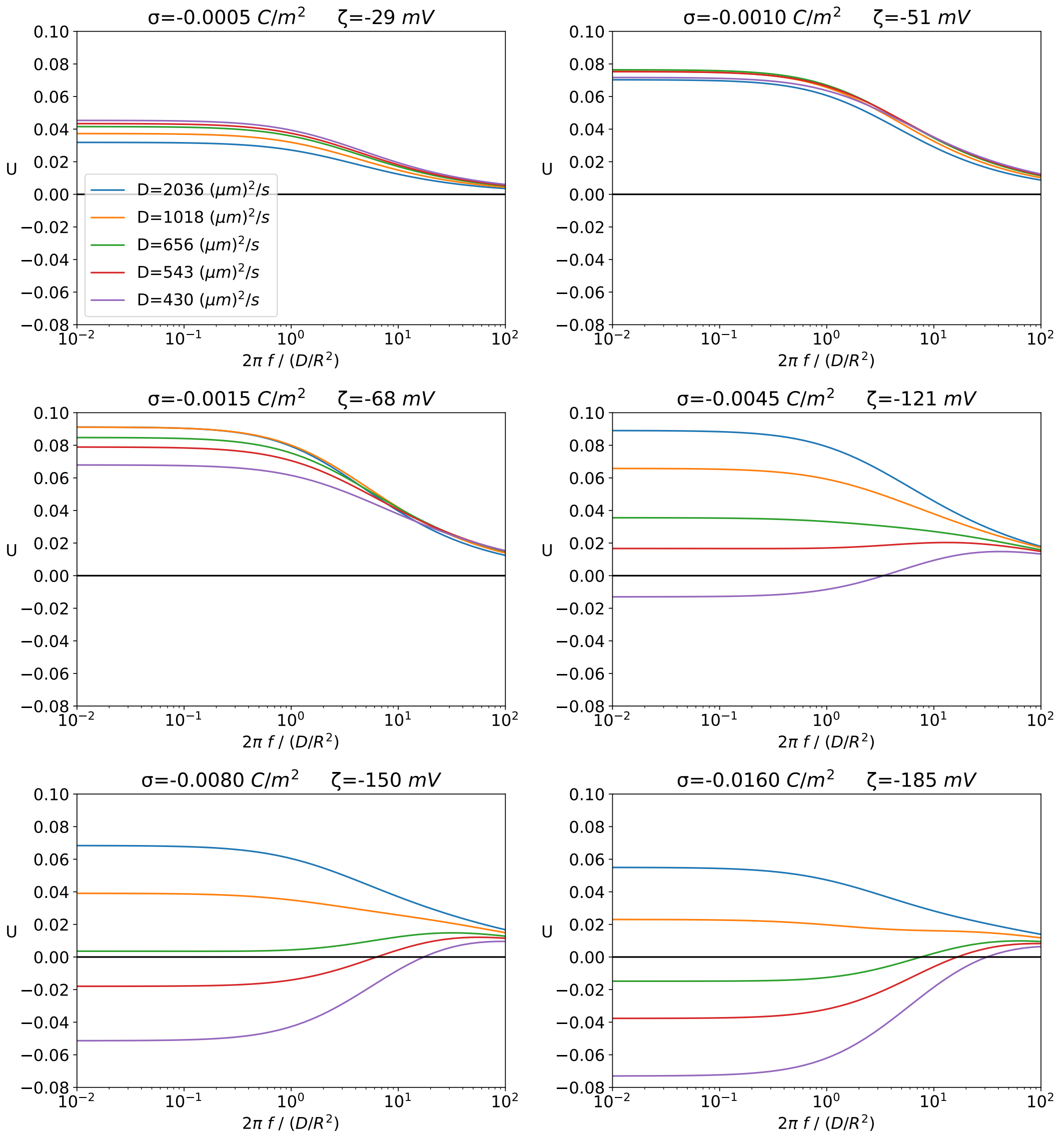


FIG. 4. Plots of the frequency response of the dimensionless flow magnitude for CPEO flows around spheres with different surface charges $\sigma$ and diffusion constants $D$. The Debye length is set to $\lambda = 43\,\mathrm{nm}$ and the particle radius is $R = 0.78\,\mu\mathrm{m}$

### IV. FUNCTIONALIZATION OF THE COLLOIDS

#### A. DNA sequences

The sequences of the oligonucleotides used for modification and cross-linking of the microparticles (in 5' to 3' direction) are listed below:

- /5AmMC6/TTCGTTTTAGTCCCATTTGTTCAGTTTTTTCAGTTTTAGCGCGGTAGTTCAGTTAGGTCA
- /5AmMC6/GTCTTTTATGCTGCTTATTCGTGTATATCCTGACCTAACTGAACTACCGCGCTAAAACTG

Here '/5AmMC6/' denotes the amino modification at the 5' end of the sequences by the oligo synthesis company (IDT).

#### B. Buffer and reagent stocks

The following buffers and reagents can be prepared in advance.

- $1\,$M MES buffer (Carl Roth: Art. No 4256.2) titrated to pH 4.8 with HCl and NaOH
- $0.1\,$M MES buffer diluted from the stock above
- $50\,$mM Borate buffer at pH 8.2 prepared by titrating boric acid (Carl Roth: Art. N0 6943.2) with HCl and NaOH
- 5'-amino-modified DNA diluted in deionized water to a concentration of $1\,$mM

Ethyl-3-(3-dimethyl-aminopropyl) carbodiimide (EDC) (Merck: Art. No. E6383-1G) is stored in dry form in small aliquots of approximately $20\,$mg at $-20\,°$C and is later diluted in deionized water just before starting the bioconjugation reaction.

#### C. Colloid concentrations

Carboxylated silica spheres with diameters $1.01\,\mu$m (Lot: $SiO_2$−COOH-AR756-5ml) and $2.12\,\mu$m (Lot: $SiO_2$−COOH-AR1060-5ml) were purchased from the microParticles GmbH. The colloids come at a weight per volume concentration of $c_W = 0.05\,$g/mL. This can be translated into a number density $c$ via

$$c = \frac{c_W}{\rho \frac{4}{3}\pi \left(\frac{d}{2}\right)^3}$$

were $\rho = 1.85\,\text{g/cm}^3$ is the density of the colloids and $d$ is their diameter. The number density of the colloids is thus given by

$$c_{1.01} = \frac{50 \cdot 10^9}{\text{mL}}$$
$$c_{2.12} = \frac{5.42 \cdot 10^9}{\text{mL}}.$$

The following relation between the colloid concentrations $c^s$ in the reaction has to be fulfilled in order to have approximately the same number of reaction sites in a sample:

$$c^s_{2.12} = \left(\frac{1.01}{2.12}\right)^2 c^s_{1.01}, \tag{7}$$

which relates the number densities via the surface area of the colloids. We choose

$$c^s_{1.01} = 10 \cdot 10^9 \frac{1}{mL}$$
$$c^s_{2.12} = 2.27 \cdot 10^9 \frac{1}{mL}.$$

as the starting point for our protocol.

#### D. Protocol

*0. Colloid start concentration*

- vortex colloid stock solutions
- $2.12\,\mu$m colloids: Pipet $419\,\mu$L of colloids and $581\,\mu$L of deionized water into a $1.5\,$mL tube (dilution 1/2.386)
- $1.01\,\mu$m colloids: Pipet $200\,\mu$L of colloids and $800\,\mu$L of deionized water into a $1.5\,$mL tube (dilution 1/5)

*1. Transfer Colloids into MES buffer*

- centrifuge the colloids at 250 rcf for 30 s and remove the supernatant (the Colloids should be sedimented)
- add 1 mL of 0.1 M MES buffer

- vortex briefly
- centrifuge the colloids at 250 rcf for 30 s (Colloids should be sedimented)
- add $100$ µL of 0.1 M MES buffer at pH 4.8
- sonicate the colloids with a ultrasound generator at maximum power settings (20 W) for 1 min

The sonication with the ultrasound generator (Bandelin: SONOPLUS UW mini20) breaks up potentially aggregated colloids.

*2. Modification reaction*

- create 1.25 M EDC stock solution: Disolve 12 mg of EDC in 50 µL deionized water
- add 50 µL of 1 mM amino-modified DNA and 10 µL of 1 M MES-buffer and 40 µL of 1.25 M EDC to a 0.5 mL tube
- vortex briefly
- add the 100 µL of the prepared colloids solution to the sample.
- vortex briefly
- incubate for 3 h on a rotator

*3. Washing procedure*

The colloids are first washed with borate buffer to neutralize the acidic MES buffer and to hydrolize unreacted activated carboxyl groups. The colloids are then washed with deionized water to remove the borate buffer and any reactants that are unspecifically adsorbed to the colloids.

- transfer the sample to a 1.5 mL tube
- add 800 µL of 50 mM borate buffer to the sample.
- repeat the following steps for 5 times

  - centrifuge at 250 rcf for 30 s (the colloids should be sedimented) and remove the supernatant
  - add 1 mL of 50 mM borate buffer to the sample
  - vortex briefly; In round 4: sonicate at max settings for 1 min and incubate for 1.5 h on a rotator

- repeat the following steps for 5 times
  - Centrifuge at 1000 rcf for 30 s (Colloids should be sedimented) and remove the supernatant
  - add 1 mL of deionized water to the sample
  - vortex briefly; in round 4: sonicate at max setting for 1 min and incubate for 1.5 h on a rotator

- sonicate the colloids with a ultrasound generator at max settings for 1 min
- create aliquots, shock freeze in liquid nitrogen and store at $-80\,^{\circ}$C

### V. RAW DATA

#### A. Buffer characterization

Tables S1, S3, S5 and S7 list all velocity measurements for all buffer conditions. Column 1 and 2 list the concentrations of the buffer ingredients. Column 3 (video no.) enumerates the videos recorded with the same buffer condition. Column 4 (dimer (µm/s)) reports the measured velocities of all dimers in a video as an ordered list. Column 5 (orientation) lists the orientations of the dimers from Column 4 encoded as 1 ('up') and -1 ('down'). Up corresponds to the case where the smaller particle of a dimer appears below the larger particle on the computer screen. Column 6 (reference (µm/s)) contains the measured velocities of all reference particles as an ordered list. Column 7 (average reference (µm/s)) lists the average velocity of the reference particles from Column 6 of one video. Column 8 (dimer corrected (µm/s)) lists the dimer velocities from Column 4 corrected by the average velocity of the reference particles from Column 7.
Tables S2, S4, S6 and S8 list the corrected velocity measurements, and the corresponding average and standard deviation for the given buffer compositions. Column 1 and 2 list the concentrations

of the buffer ingredients. Column 3 (dimer corrected (µm/s)) list all corrected dimer velocities of a given buffer composition. Column 4 (average (µm/s)) reports the average of the corrected dimer velocities. Column 5 (standard deviation (µm/s)) lists the standard deviation of the corrected dimer velocities.

TABLE I. NaOH raw data

| NaOH (µM) | $MgCl_2$ (µM) | video no. | dimer (µm/s) | orientation | reference (µm/s) | average reference (µm/s) | dimer corrected (µm/s) |
|---|---|---|---|---|---|---|---|
| 10 | 5.2 | 1 | -2.1584; 2.0315; 1.7996 | 1; -1; -1 | -0.11963 | -0.11963 | 2.0388; 2.1512; 1.9192 |
| 10 | 5.2 | 2 | -2.11; 1.8049 | 1; -1 | 0.09221 | 0.09221 | 2.2022; 1.7127 |
| 10 | 5.2 | 3 | 1.7057; 1.6759 | -1; -1 | -0.22243; -0.26056; -0.17912 | -0.2207 | 1.9264; 1.8966 |
| 25 | 0.2 | 1 | -0.62871; -0.66528 | -1; -1 | -0.16556; -0.23086; -0.21944 | -0.20529 | -0.42343; -0.45999 |
| 25 | 0.2 | 2 | -0.22598 | 1 | -0.33294; -0.11588 | -0.22441 | 0.0015664 |
| 25 | 0.2 | 3 | -0.32598 | -1 | -0.1046 | -0.1046 | -0.22138 |
| 25 | 5.2 | 1 | 3.081; 2.9794 | -1; -1 | 0.016034; 0.10796; 0.10584 | 0.076612 | 3.0044; 2.9028 |
| 25 | 5.2 | 2 | -3.098; 3.1878 | 1; -1 | -0.033192 | -0.033192 | 3.0648; 3.221 |
| 25 | 5.2 | 3 | -3.8004; 3.4204 | 1; -1 | -0.24992; -0.36162; -0.092779; 0.1997 | -0.12616 | 3.6742; 3.5466 |
| 50 | 0.2 | 1 | -0.69124; 1.0838; 0.81608 | -1; 1; 1 | 0.15563 | 0.15563 | -0.84687; -0.92813; -0.66045 |
| 50 | 0.2 | 2 | -1.0354; -1.1525 | -1; -1 | 0.065881 | 0.065881 | -1.1013; -1.2184 |
| 50 | 0.2 | 3 | -0.69959; -0.90219; -0.88047 | -1; -1; -1 | -0.12209 | -0.12209 | -0.57749; -0.78009; -0.75838 |
| 50 | 5.2 | 1 | 0.36514; -0.75861 | -1; 1 | -0.011512 | -0.011512 | 0.37665; 0.7471 |
| 50 | 5.2 | 2 | 0.49612; -0.9068 | -1; 1 | -0.12991; 0.10312; 0.039071 | 0.0040941 | 0.49203; 0.9109 |
| 50 | 5.2 | 3 | -0.71939; -0.87973 | 1; 1 | -0.09121 | -0.09121 | 0.62818; 0.78852 |
| 100 | 5.2 | 1 | -0.31532; -0.17464 | 1; -1 | 0.049767 | 0.049767 | 0.36509; -0.22441 |
| 100 | 5.2 | 2 | 0.05744; -0.03439 | 1; 1 | -0.16706; 0.050236; -0.1582 | -0.091673 | -0.14911; -0.057283 |
| 100 | 5.2 | 3 | -0.044189; 0.29934 | 1; 1 | 0.01613; -0.020345; -0.1041 | -0.036106 | 0.0080835; -0.33544 |

TABLE II. NaOH corrected velocities

| NaOH (µM) | $MgCl_2$ (µM) | dimer corrected (µm/s) | average (µm/s) | standard deviation (µm/s) |
|---|---|---|---|---|
| 10 | 5.2 | 2.0388; 2.1512; 1.9192; 2.2022; 1.7127; 1.9264; 1.8966 | 1.9782 | 0.16687 |
| 25 | 0.2 | -0.42343; -0.45999; 0.0015664; -0.22138 | -0.27581 | 0.21261 |
| 25 | 5.2 | 3.0044; 2.9028; 3.0648; 3.221; 3.6742; 3.5466 | 3.2356 | 0.31069 |
| 50 | 0.2 | -0.84687; -0.92813; -0.66045; -1.1013; -1.2184; -0.57749; -0.78009; -0.75838 | -0.85889 | 0.2164 |
| 50 | 5.2 | 0.37665; 0.7471; 0.49203; 0.9109; 0.62818; 0.78852 | 0.65723 | 0.19825 |
| 100 | 5.2 | 0.36509; -0.22441; -0.14911; -0.057283; 0.0080835; -0.33544 | -0.065514 | 0.24331 |

### B. Electric field strength and frequency characterization

Table S9, S10 and S11 each contain data obtained from measurements made on one individual dimer and a reference particle. Column 1 and 2 list the applied electric field amplitude and frequency. Column 3 lists the measured velocity of the dimer. Column 4 lists the velocity of the reference particle. Column 5 lists the corrected velocity of the dimer.

TABLE III. Tris raw data

| Tris (μM) | $MgCl_2$ (μM) | video no. | dimer (μm/s) | orientation | reference (μm/s) | average reference (μm/s) | dimer corrected (μm/s) |
|---|---|---|---|---|---|---|---|
| 10 | 5.2 | 1 | 2.1867; 2.112 | -1; -1 | -0.011658 | -0.011658 | 2.1983; 2.1237 |
| 10 | 5.2 | 2 | 2.3178 | -1 | -0.28296; -0.093157 | -0.18806 | 2.5058 |
| 10 | 5.2 | 3 | 2.0876; -2.5115 | -1; 1 | -0.01149 | -0.01149 | 2.099; 2.5 |
| 10 | 5.2 | 4 | 2.384; -2.5342 | -1; 1 | -0.31553; 0.031664 | -0.14193 | 2.526; 2.3922 |
| 25 | 5.2 | 1 | 2.7714; -1.5856 | -1; 1 | 0.035405 | 0.035405 | 2.736; 1.621 |
| 25 | 5.2 | 2 | -2.8863; 2.7368 | 1; -1 | 0.0074493 | 0.0074493 | 2.8938; 2.7293 |
| 25 | 5.2 | 3 | -2.1166; -2.9565; 2.956 | 1; 1; -1 | -0.0031657 | -0.0031657 | 2.1135; 2.9533; 2.9592 |
| 50 | 0.2 | 1 | 0.24035; -0.29358; -0.29227 | -1; 1; 1 | -0.07472 | -0.07472 | 0.31507; 0.21886; 0.21755 |
| 50 | 0.2 | 2 | -0.18728; 0.3081; 0.067422 | 1; -1; -1 | -0.10733; 0.014156 | -0.046585 | 0.1407; 0.35468; 0.11401 |
| 50 | 0.2 | 3 | -0.41175; -0.35746 | 1; 1 | 0.081775 | 0.081775 | 0.49352; 0.43924 |
| 50 | 5.2 | 1 | 2.2018; -2.4064 | -1; 1 | -0.082894 | -0.082894 | 2.2847; 2.3235 |
| 50 | 5.2 | 2 | 2.2391; -2.415 | -1; 1 | 0.042677; -0.078361 | -0.017842 | 2.257; 2.3971 |
| 50 | 5.2 | 3 | 2.4588; -2.2549 | -1; 1 | -0.04447 | -0.04447 | 2.5033; 2.2105 |
| 100 | 5.2 | 1 | 1.1616; -1.736; -1.8208 | -1; 1; 1 | -0.22214; -0.30256 | -0.26235 | 1.424; 1.4736; 1.5584 |
| 100 | 5.2 | 2 | 1.7564; -1.4701 | -1; 1 | 0.083002; 0.033984 | 0.058493 | 1.6979; 1.5286 |
| 100 | 5.2 | 3 | -1.2916; 1.5386; -1.7132 | 1; -1; 1 | -0.071143; 0.016444 | -0.027349 | 1.2642; 1.566; 1.6859 |
| 250 | 5.2 | 1 | 0.82139; -0.63782 | -1; 1 | -0.016064 | -0.016064 | 0.83745; 0.62176 |
| 250 | 5.2 | 2 | -0.87382; -0.94419 | 1; 1 | -0.03497 | -0.03497 | 0.83885; 0.90922 |
| 250 | 5.2 | 3 | -0.95719; -0.82211; 0.80946 | 1; 1; -1 | -0.13714; 0.20802 | 0.035439 | 0.99263; 0.85755; 0.77402 |
| 500 | 5.2 | 1 | -0.55723; 0.51097; -0.45549 | 1; -1; 1 | -0.065543 | -0.065543 | 0.49169; 0.57651; 0.38994 |
| 500 | 5.2 | 3 | -0.54143; 0.78223 | 1; -1 | -0.2473 | -0.2473 | 0.29413; 1.0295 |

TABLE IV. Tris corrected velocities

| Tris (μM) | $MgCl_2$ (μM) | dimer corrected (μm/s) | average (μm/s) | standard deviation (μm/s) |
|---|---|---|---|---|
| 10 | 5.2 | 2.1983; 2.1237; 2.5058; 2.099; 2.5; 2.526; 2.3922 | 2.335 | 0.18938 |
| 25 | 5.2 | 2.736; 1.621; 2.8938; 2.7293; 2.1135; 2.9533; 2.9592 | 2.5723 | 0.51075 |
| 50 | 0.2 | 0.31507; 0.21886; 0.21755; 0.1407; 0.35468; 0.11401; 0.49352; 0.43924 | 0.2867 | 0.13732 |
| 50 | 5.2 | 2.2847; 2.3235; 2.257; 2.3971; 2.5033; 2.2105 | 2.3293 | 0.10607 |
| 100 | 5.2 | 1.424; 1.4736; 1.5584; 1.6979; 1.5286; 1.2642; 1.566; 1.6859 | 1.5248 | 0.14114 |
| 250 | 5.2 | 0.83745; 0.62176; 0.83885; 0.90922; 0.99263; 0.85755; 0.77402 | 0.83307 | 0.11551 |
| 500 | 5.2 | 0.49169; 0.57651; 0.38994; 0.29413; 1.0295 | 0.55636 | 0.28501 |

TABLE V. NaCl raw data

| NaCl (μM) | $MgCl_2$ (μM) | video no. | dimer (μm/s) | orientation | reference (μm/s) | average reference (μm/s) | dimer corrected (μm/s) |
|---|---|---|---|---|---|---|---|
| 25 | 5.2 | 1 | 0.2565; -0.090399 | -1; -1 | -0.047559; -0.10738 | -0.077468 | 0.33397; -0.01293 |
| 25 | 5.2 | 2 | -0.0010654; -0.072518 | 1; -1 | -0.083108 | -0.083108 | -0.082042; 0.01059 |
| 25 | 5.2 | 3 | -0.058622; 0.077811; 0.20893 | 1; -1; -1 | -0.0033616 | -0.0033616 | 0.055261; 0.081173; 0.21229 |
| 50 | 0.2 | 1 | -0.07064 | -1 | -0.10322 | -0.10322 | 0.032583 |
| 50 | 0.2 | 2 | -0.30739; 0.14427 | -1; 1 | -0.20242 | -0.20242 | -0.10497; -0.34669 |
| 50 | 0.2 | 3 | 0.23731; -0.0034925 | 1; 1 | 0.022871; 0.14171 | 0.08229 | -0.15502; 0.085783 |
| 50 | 5.2 | 1 | 0.18548; -0.60578 | -1; 1 | 0.10408; -0.16269; -0.13499 | -0.064532 | 0.25002; 0.54125 |
| 50 | 5.2 | 2 | 0.35203; -0.4395 | -1; 1 | 0.013808; 0.12143 | 0.067621 | 0.28441; 0.50712 |
| 50 | 5.2 | 3 | -0.36842; 0.49881 | 1; -1 | 0.041369; -0.13753 | -0.048079 | 0.32034; 0.54689 |
| 50 | 5.2 | 4 | -0.62647; 0.5147; -0.6237 | 1; -1; 1 | -0.011805; 0.0073405 | -0.0022323 | 0.62424; 0.51693; 0.62146 |
| 100 | 0.2 | 1 | -0.3359; -0.070846; -0.32053 | 1; 1; 1 | -0.39082; -0.018323 | -0.20457 | 0.13133; -0.13372; 0.11596 |
| 100 | 0.2 | 2 | -0.050905 | -1 | -0.15048; -0.098426; -0.10022; 0.21803; -0.13712 | -0.053645 | 0.0027406 |
| 100 | 0.2 | 3 | 0.041658; 0.27696; 0.091807 | -1; -1; -1 | 0.035803; 0.092218 | 0.06401 | -0.022352; 0.21295; 0.027797 |
| 100 | 0.2 | 4 | -0.19775 | 1 | -0.035193; 0.050215; -0.069265 | -0.018081 | 0.17967 |
| 100 | 5.2 | 1 | 0.54022; -0.76891 | -1; 1 | -0.064309 | -0.064309 | 0.60453; 0.7046 |
| 100 | 5.2 | 2 | -0.9319; -0.94887 | 1; 1 | -0.061837; 0.0075117; -0.099579 | -0.051301 | 0.88059; 0.89757 |
| 100 | 5.2 | 3 | 0.51296; 0.58286 | -1; -1 | 0.1632 | 0.1632 | 0.34976; 0.41966 |
| 100 | 5.2 | 4 | 0.59116; 0.54284 | -1; -1 | -0.0032619 | -0.0032619 | 0.59442; 0.5461 |
| 250 | 5.2 | 1 | 0.35226; -0.81461 | -1; 1 | 0.10234 | 0.10234 | 0.24992; 0.91694 |
| 250 | 5.2 | 2 | 0.57366; 0.52717; -0.74718 | -1; -1; 1 | -0.012219 | -0.012219 | 0.58588; 0.53939; 0.73496 |
| 250 | 5.2 | 3 | -0.43561; 0.52548 | 1; -1 | -0.007625 | -0.007625 | 0.42798; 0.53311 |
| 500 | 5.2 | 1 | -0.23991; 0.53692; -0.27462 | 1; -1; 1 | 0.118; -0.19101 | -0.036507 | 0.20341; 0.57343; 0.23812 |
| 500 | 5.2 | 2 | 0.69386; -0.13413 | -1; 1 | -0.16074 | -0.16074 | 0.8546; -0.026605 |
| 500 | 5.2 | 3 | 0.36688; 0.44444 | -1; -1 | -0.10812 | -0.10812 | 0.475; 0.55257 |

TABLE VI. NaCl corrected velocities

| NaCl (μM) | $MgCl_2$ (μM) | dimer corrected (μm/s) | average (μm/s) | standard deviation (μm/s) |
|---|---|---|---|---|
| 25 | 5.2 | 0.33397; -0.01293; -0.082042; 0.01059; 0.055261; 0.081173; 0.21229 | 0.085472 | 0.14266 |
| 50 | 0.2 | 0.032583; -0.10497; -0.34669; -0.15502; 0.085783 | -0.097663 | 0.17027 |
| 50 | 5.2 | 0.25002; 0.54125; 0.28441; 0.50712; 0.32034; 0.54689; 0.62424; 0.51693; 0.62146 | 0.46807 | 0.14427 |
| 100 | 0.2 | 0.13133; -0.13372; 0.11596; 0.0027406; -0.022352; 0.21295; 0.027797; 0.17967 | 0.064296 | 0.11614 |
| 100 | 5.2 | 0.60453; 0.7046; 0.88059; 0.89757; 0.34976; 0.41966; 0.59442; 0.5461 | 0.62465 | 0.19693 |
| 250 | 5.2 | 0.24992; 0.91694; 0.58588; 0.53939; 0.73496; 0.42798; 0.53311 | 0.56974 | 0.21321 |
| 500 | 5.2 | 0.20341; 0.57343; 0.23812; 0.8546; -0.026605; 0.475; 0.55257 | 0.41007 | 0.29205 |

### TABLE VII. $MgCl_2$ raw data

| $MgCl_2$ (μM) | video no. | dimer (μm/s) | orientation | reference (μm/s) | average reference (μm/s) | dimer corrected (μm/s) |
|---|---|---|---|---|---|---|
| 5 | 4 | -0.1007 | 1 | 0.12329; 0.078374; 0.08571 | 0.095791 | 0.19649 |
| 5 | 1 | 0.89903; -0.73302 | 1; -1 | 0.033914 | 0.033914 | -0.86511; -0.76694 |
| 5 | 2 | 0.50023; -0.76023 | 1; -1 | 0.040048 | 0.040048 | -0.46019; -0.80028 |
| 5 | 3 | 0.61077; 0.64962 | 1; 1 | -0.071589 | -0.071589 | -0.68236; -0.72121 |
| 10 | 1 | -0.77615 | -1 | -0.11222 | -0.11222 | -0.66393 |
| 10 | 2 | 0.81232 | 1 | -0.049807; 0.027225; -0.089389; 0.061394 | -0.012644 | -0.82496 |
| 10 | 3 | -0.67114; -0.61298 | -1; -1 | 0.16781; 0.0054852; -0.045543; 0.050324 | 0.04452 | -0.71566; -0.6575 |
| 10 | 4 | 0.94059 | 1 | -0.1349 | -0.1349 | -1.0755 |
| 10 | 5 | -0.44429 | -1 | -0.061833; 0.035126; 0.065325 | 0.012873 | -0.45717 |
| 25 | 1 | -0.2533; 0.49167 | 1; 1 | -0.20812 | -0.20812 | 0.045177; -0.69979 |
| 25 | 2 | -0.68736; -0.14944 | -1; -1 | -0.041324; -0.064957; -0.036963 | -0.047748 | -0.63961; -0.10169 |
| 25 | 3 | -0.40842 | -1 | -0.099255 | -0.099255 | -0.30917 |
| 25 | 4 | -0.31028; 0.63511 | -1; 1 | -0.021058 | -0.021058 | -0.28922; -0.65617 |
| 50 | 1 | -0.043009 | -1 | 0.0075411 | 0.0075411 | -0.05055 |
| 50 | 2 | -0.1422; -0.13616 | -1; 1 | 0.0054712 | 0.0054712 | -0.14768; 0.14163 |
| 50 | 3 | -0.079519; 0.080918 | -1; 1 | -0.045867 | -0.045867 | -0.033653; -0.12679 |
| 100 | 1 | 0.11921 | -1 | 0.0010629 | 0.0010629 | 0.11815 |
| 100 | 2 | 0.18407 | -1 | 0.074598; 0.051411 | 0.063005 | 0.12106 |
| 100 | 3 | 0.31911; 0.23815 | -1; -1 | -0.0078792; -0.064006 | -0.035943 | 0.35505; 0.27409 |
| 100 | 4 | -0.46523 | 1 | 0.066031; 0.24479 | 0.15541 | 0.62064 |
| 100 | 5 | 0.29586; -0.21409 | -1; 1 | -0.060438 | -0.060438 | 0.35629; 0.15366 |
| 250 | 1 | -0.2334 | 1 | -0.062887 | -0.062887 | 0.17051 |
| 250 | 2 | 0.24637 | -1 | 0.0055673; -0.030459; -0.0033249 | -0.0094054 | 0.25577 |
| 250 | 3 | 0.33192; -0.28766 | -1; 1 | 0.061578 | 0.061578 | 0.27034; 0.34924 |

### TABLE VIII. $MgCl_2$ corrected velocities

| $MgCl_2$ (μM) | dimer corrected (μm/s) | average (μm/s) | standard deviation (μm/s) |
|---|---|---|---|
| 5 | -0.86511; -0.76694; -0.46019; -0.80028; -0.68236; -0.72121 | -0.71601 | 0.14038 |
| 10 | -0.66393; -0.82496; -0.71566; -0.6575; -1.0755; -0.45717 | -0.73245 | 0.20621 |
| 25 | 0.045177; -0.69979; -0.63961; -0.10169; -0.30917; -0.28922; -0.65617 | -0.37864 | 0.29366 |
| 50 | -0.05055; -0.14768; 0.14163; -0.033653; -0.12679; 0.19649 | -0.0034233 | 0.14154 |
| 100 | 0.11815; 0.12106; 0.35505; 0.27409; 0.62064; 0.35629; 0.15366 | 0.28556 | 0.18015 |
| 250 | 0.17051; 0.25577; 0.27034; 0.34924 | 0.26147 | 0.073229 |

### TABLE IX. Swimmer 1

| electric field (mV/μm) | frequency (Hz) | dimer (μm/s) | reference (μm/s) | dimer corrected (μm/s) |
|---:|---:|---:|---:|---:|
| 3.05 | 250 | -0.04425 | -0.05515 | 0.01089 |
| 4.57 | 250 | 0.18415 | -0.07716 | 0.26130 |
| 6.09 | 250 | -0.36709 | -0.05731 | 0.30978 |
| 7.61 | 250 | -0.58703 | -0.06520 | 0.52183 |
| 9.14 | 250 | 0.73478 | -0.10905 | 0.84382 |
| 10.66 | 250 | -1.01412 | -0.04195 | 0.97218 |
| 12.18 | 250 | 1.08429 | -0.07286 | 1.15715 |
| 13.71 | 250 | -1.58886 | -0.00330 | 1.58556 |
| 15.23 | 250 | 1.90124 | 0.06676 | 1.83448 |
| 16.75 | 250 | -2.27900 | 0.02538 | 2.30438 |
| 16.75 | 500 | -1.90459 | 0.10462 | 2.00922 |
| 16.75 | 750 | 1.27442 | 0.04976 | 1.22466 |
| 16.75 | 1000 | 0.84186 | -0.06135 | 0.90321 |
| 16.75 | 1500 | -0.14115 | 0.23458 | 0.37573 |
| 16.75 | 2000 | 0.00711 | 0.11272 | 0.10561 |

TABLE X. Swimmer 2

| electric field (mV/μm) | frequency (Hz) | dimer (μm/s) | reference (μm/s) | dimer corrected (μm/s) |
|---:|---:|---:|---:|---:|
| 3.05 | 250 | -0.07672 | -0.03346 | 0.04326 |
| 4.57 | 250 | -0.23488 | -0.05509 | 0.17980 |
| 6.09 | 250 | 0.29167 | -0.13350 | 0.42517 |
| 7.61 | 250 | 0.50567 | -0.10702 | 0.61269 |
| 9.14 | 250 | -0.96007 | -0.15874 | 0.80133 |
| 10.66 | 250 | 0.91369 | -0.09650 | 1.01020 |
| 12.18 | 250 | 1.37226 | -0.08227 | 1.45453 |
| 13.71 | 250 | -1.85958 | -0.19601 | 1.66357 |
| 15.23 | 250 | 2.01217 | -0.13712 | 2.14930 |
| 16.75 | 250 | -2.41953 | 0.00508 | 2.42461 |
| 16.75 | 500 | 1.90613 | -0.27114 | 2.17727 |
| 16.75 | 750 | -1.64252 | -0.34347 | 1.29905 |
| 16.75 | 1000 | -1.02669 | -0.22037 | 0.80632 |
| 16.75 | 1500 | -0.51470 | -0.02538 | 0.48932 |
| 16.75 | 2000 | -0.40936 | -0.36060 | 0.04876 |

TABLE XI. Swimmer 3

| electric field (mV/μm) | frequency (Hz) | dimer (μm/s) | reference (μm/s) | dimer corrected (μm/s) |
|---:|---:|---:|---:|---:|
| 3.05 | 250 | -0.05914 | 0.00575 | 0.06489 |
| 4.57 | 250 | 0.13688 | -0.03549 | 0.17237 |
| 6.09 | 250 | 0.16393 | -0.06861 | 0.23254 |
| 7.61 | 250 | 0.37440 | -0.14963 | 0.52403 |
| 9.14 | 250 | 0.53618 | -0.12732 | 0.66351 |
| 10.66 | 250 | -1.00302 | -0.16564 | 0.83737 |
| 12.18 | 250 | -1.13091 | 0.05384 | 1.18474 |
| 13.71 | 250 | 1.15126 | -0.26317 | 1.41444 |
| 15.23 | 250 | 1.65991 | -0.01931 | 1.67923 |
| 16.75 | 250 | -1.95483 | -0.12504 | 1.82979 |
| 16.75 | 500 | -1.72144 | -0.12492 | 1.59652 |
| 16.75 | 750 | -1.33342 | -0.13304 | 1.20038 |
| 16.75 | 1000 | 0.71604 | -0.08498 | 0.80102 |
| 16.75 | 1500 | -0.41806 | -0.11272 | 0.30534 |
| 16.75 | 2000 | 0.17822 | 0.09728 | 0.08093 |

[1] Fernández-Mateo, R., García-Sánchez, P., Calero, V., Morgan, H. & Ramos, A. Stationary electro-osmotic flow driven by ac fields around charged dielectric spheres. *Journal of Fluid Mechanics* **924** (2021). URL https://www.cambridge.org/core/journals/journal-of-fluid-mechanics/article/stationary-electroosmotic-flow-driven-by-ac-fields-around-charged-dielectric-spheres/4D37A1F8B31B7712D0855E0F9E6DCF54.

[2] Khair, A. S. & Balu, B. Breaking electrolyte symmetry in induced-charge electro-osmosis. *Journal of Fluid Mechanics* **905** (2020).

[3] Squires, T. M. & Bazant, M. Z. Induced-charge electro-osmosis. *Journal of Fluid Mechanics* **509**, 217–252 (2004).

[4] Schnitzer, O. & Yariv, E. Strong electro-osmotic flows about dielectric surfaces of zero surface charge. *Physical review. E, Statistical, nonlinear, and soft matter physics* **89**, 043005 (2014).
